# Primary auditory cortex is necessary for the acquisition and expression of categorical behavior

**DOI:** 10.1101/2024.02.02.578700

**Authors:** Rebecca F. Krall, Rachel M. Cassidy, Madan Ghimire, Callista N. Chambers, Megan P. Arnold, Lauren I. Brougher, Justin Chen, Rishi Deshmukh, Hailey B. King, Harry J. Morford, John M. Wiemann, Ross S. Williamson

## Abstract

The primary auditory cortex (ACtx) is critically involved in the association of sensory information with specific behavioral outcomes. Such sensory-guided behaviors are necessarily brain-wide endeavors, requiring a plethora of distinct brain areas, including those that are involved in aspects of decision making, motor planning, motor initiation, and reward prediction. ACtx comprises a number of distinct excitatory cell-types that allow for the brain-wide propagation of behaviorally-relevant sensory information. Exactly how ACtx involvement changes as a function of learning, as well as the functional role of distinct excitatory cell-types is unclear. Here, we addressed these questions by designing a two-choice auditory task in which water-restricted, head-fixed mice were trained to categorize the temporal rate of a sinusoidal amplitude modulated (sAM) noise burst and used transient cell-type specific optogenetics to probe ACtx necessity across the duration of learning. Our data demonstrate that ACtx is necessary for the ability to categorize the rate of sAM noise, and this necessity remains stable across learning. ACtx silencing substantially altered the behavioral strategies used to solve the task by introducing a fluctuating choice bias and increasing dependence on prior decisions. Furthermore, ACtx silencing did not impact the animal’s motor report, suggesting that ACtx is necessary for the conversion of sensation to action. Targeted inhibition of both intratelencephalic and extratelencephalic projections on just 20% of trials had a modest effect on task performance, but significantly degraded learning. Taken together, our data shows that ACtx plays a critical role in both the acquisition and expression of categorical behavior, and that distinct excitatory projections play important roles in learning and plasticity.

## Introduction

Effectively engaging with the world around us requires intricate perceptual processing that enables us to interpret the sensory environment and react accordingly within the given context. The learning process of associating sensory information with specific behavioral outcomes is essential for the rapid selection of optimal motor actions and often depends on the categorization of stimuli based on their expected outcome [1, 2]. Categorization is a computational process that involves a complex transformation of sensory information into discrete perceptual categories [3–5]. Although much work has implicated multiple sensory cortices as being involved in different aspects of categorization (and sensory-guided behavior in general) [6–9], the role of sensory cortex in both the acquisition and expression of categorical behavior is unclear.

Learning to categorize auditory stimuli based on their perceptual or behavioral relevance can induce dynamic changes in auditory cortical activity. During the acquisition phase of auditory category learning, both animal and human studies consistently demonstrate that neural responses shift to reflect the behavioral relevance of acoustic cues. Rather than passively encoding acoustic properties, cortical activity becomes increasingly aligned with category structure, reflecting a transformation from sensory-driven to behaviorally relevant representations. Such categorical reorganization has been observed across species, including in gerbils, where the emergence of categorization coincides with abrupt transitions in spatiotemporal cortical dynamics [10, 11]; in mice, where auditory cortical ensembles reorganize during learning of tone categories [8]; in non-human primates, where neurons develop categorical tuning to speech sounds [2]; and in humans, where category learning reshapes distributed cortical representations [12, 13]. Computational modeling supports the view that such category-related plasticity may arise from enhanced representation of relevant stimuli and recruitment of discriminative ensemble codes [14, 15]. However, these studies have primarily emphasized frequency-based stimuli and much less is known about how ACtx supports learning of categories based on temporal features, such as amplitude modulation rate, which impose additional demands on temporal integration and may engage distinct cortical mechanisms for temporal coding and perceptual abstraction.

Once an auditory category has been acquired, its expression during behavior is thought to rely on a stable yet flexible cortical code that supports perceptual decisions. Activity in ACtx is modulated by context and task demands, and population codes in ACtx can reflect not just stimulus properties, but also internal variables such as expectation, attention, or decision outcomes [11, 16]. Yet, the degree to which expression relies on ACtx is not fully established. While many studies have reported that primary auditory cortex (ACtx) inactivation can impair or completely abolish auditory-guided behavior [17–22], several studies have shown that ACtx inactivation can lead to little or no effect [23–25]. Although there exists significant heterogeneity in task conditions, inactivation method, and animal model [26], most evidence points towards ACtx necessity depending critically on both stimulus choice and complexity of task parameters [27]. More complex auditory objects, especially those that require temporal integration, are more likely to necessitate ACtx involvement. Such involvement is less likely to be critical for simple feature discrimination. Exactly how such ACtx involvement changes as a function of learning represents a significant gap in knowledge.

Learning to respond accurately to incoming sound stimuli is necessarily a brain-wide endeavour, requiring a plethora of distinct brain areas, including those that are involved in aspects of decision making, motor planning, motor initiation, and reward prediction [28–31]. Indeed, ACtx is comprised of distinct excitatory cell-types that facilitate the flow of sensory information to many downstream intra- and extratelencephalic targets in parietal and frontal cortices, as well as in the midbrain, thalamus, and striatum [32–37]. Although several studies have suggested distinct behavioral roles for excitatory projections (across multiple sensory modalities) [35, 38–45], few studies to date have assayed their specific contributions to the acquisition and expression of a sensory-guided behavior using a standardized task.

To investigate the role of ACtx in the acquisition and expression of categorical auditory behavior, we developed a head-fixed categorization task (adapted from [8]) in which mice learned to categorize temporally modulated sounds based on their sinusoidal amplitude modulation (sAM) rate. Throughout the course of learning, we used temporally precise, cell-type-specific optogenetic inhibition to transiently silence excitatory ACtx neurons. We found that acute ACtx inactivation significantly impaired behavioral performance, indicating that cortical activity is essential for the formation and stabilization of categorical representations. In addition to reducing psychometric sensitivity, ACtx inactivation profoundly altered the behavioral strategies used to perform the task, inducing a fluctuating choice bias and increasing dependence of prior decisions. Importantly, we show that ACtx inactivation does not impact motor reports, indicating that ACtx is essential not for executing learned actions per se, but for transforming sensory input into task-relevant categorical judgments. Finally, we show that targeted inactivation of both intratelencephalic and extratelencephalic projections on just 20% of trials has a modest effect on task performance, but significantly degrades learning. Collectively, these findings demonstrate that ACtx is required for both the acquisition and expression of categorical behavior, and that distinct excitatory projection neuron subtypes play important roles in learning and plasticity.

## Results

### ACtx is necessary for accurate categorization of sAM noise

To investigate the involvement of ACtx in categorical decision making, we designed a two-choice auditory categorization task in which water-restricted, head-fixed mice were trained to categorize the rate of a sinusoidal amplitude modulated (sAM) noise burst (**Figure 1A**). Mice were tasked with categorizing sAM noise as either “slow” or “fast” by directionally licking left or right spouts to earn a small water reward. The stimuli consisted of sAM rates spanning 2 - 32 Hz in 0.5 octave increments, presented at 70 dB SPL, including an ”uncategorizable“ 8 Hz stimulus at the category boundary that was rewarded probabilistically (50% left, 50% right). Inter-trial intervals were exponentially distributed, and ranged from 4 to 7 s. Mice were initially introduced to an easy version of the task in which they had to categorize only the two slowest (2, 2.4 Hz) and two fastest sounds (22.8, 32 Hz), allowing them to learn the association between sAM rate and spout. To facilitate the inhibition of pan-excitatory neural activity we selectively expressed a Soma-targeted *Guillardia theta* anion-conducting channelrhodopsin (stGtACR2) in CAMKII-expressing neurons within ACtx [46]. The stGtACR2 was activated via blue light (1.1 sec, 2 mW) delivered through an optic fiber, initiated 100 ms prior to sound onset and lasted for the duration of the sound (**Figure 1B**). Optogenetic inhibition was introduced in a pseudo-random 20% of trials once mice demonstrated performance above chance levels in the easy task (*>*60% correct over 200 easy trials). Mice were considered proficient at the easy task when they achieved 85% accuracy over 200 trials (**Figure 1C**). Subsequently, all additional sAM rates (4 - 16 Hz) were introduced and mice were trained to reach an expert level, defined as accuracy above 75% correct across “hard” sounds (4, 5.6, 11.2, 16 Hz). Mice readily learned this task. Mice achieved the threshold for optogenetics within 2.8 ± 0.2 sessions (925.6 ± 66.7 trials), achieved the proficient threshold in 8.8 ± 0.5 sessions (3558.3 ± 237.3 trials) and achieved the expert threshold in 13.4 ± 0.5 sessions (5591.2 ± 306.6 trials). On average, mice completed 430.1 ± 23.6 trials per session.

**Figure 1:**
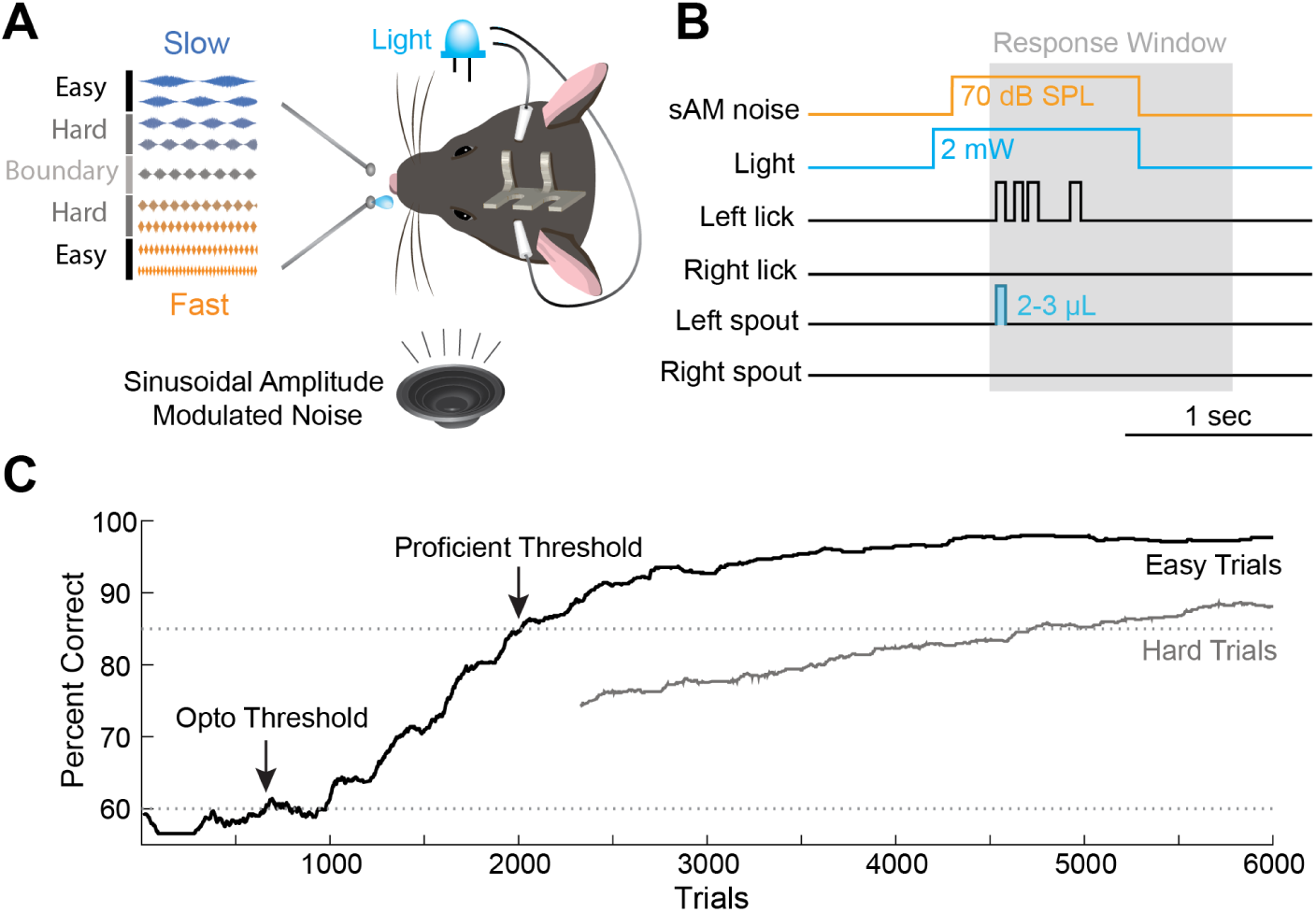
sAM Categorization Task. **A**: Schematic representation of the stimulus set and behavioral setup. Head-fixed mice are trained to categorize sAM noise as “slow” or “fast”, based on the rate of sinusoidal modulation. Blue light is delivered on 20% of trials via optic fibers positioned over the ACtx. **B**: Schematic illustration of the trial structure, outlining the sequence of events within a single behavioral trial, from the presentation of sAM noise to the mouse’s behavioral response, and details the timing of the optogenetic inhibition relative to the sound onset. **C**: Exemplar task performance across multiple trials for a single mouse. Performance is visualized as a rolling accuracy calculated over a 200 trial window. Thresholds for optogenetic introduction and task proficiency are denoted by horizontal dotted lines at 60% and 85%, respectively.

To verify the efficacy of our optogenetic silencing approach, we performed extracellular recordings (N=2) using a high-density, 64 channel, silicon probe to sample neural activity across the full depth of the ACtx (**Figure 2A**). Optogenetic activation of stGtACR2 in excitatory neurons produced near-complete suppression of spiking activity throughout the ACtx column, effectively abolishing both spontaneous and sound-evoked firing (**Figure 2B**). To quantify the extent of optogenetic-induced modulation, we computed an optogenetic modulation index for each neuron (-1 indicates complete suppression) in response to the same sAM stimuli used in the behavioral task. Of 33 sound-responsive and 10 non-responsive neurons, 29 neurons were completely silenced (67%), with a MI of -1 (**Figure 2C, top**). A parallel analysis of spontaneous activity revealed a comparable degree of silencing (**Figure 2C, bottom**). In both cases, almost 90% of neurons suppressed their firing rates by more than 90%. These results confirm that stGtACR2 provides robust and widespread inhibition across cortical layers, sufficient to functionally silence the ACtx during task engagement.

**Figure 2:**
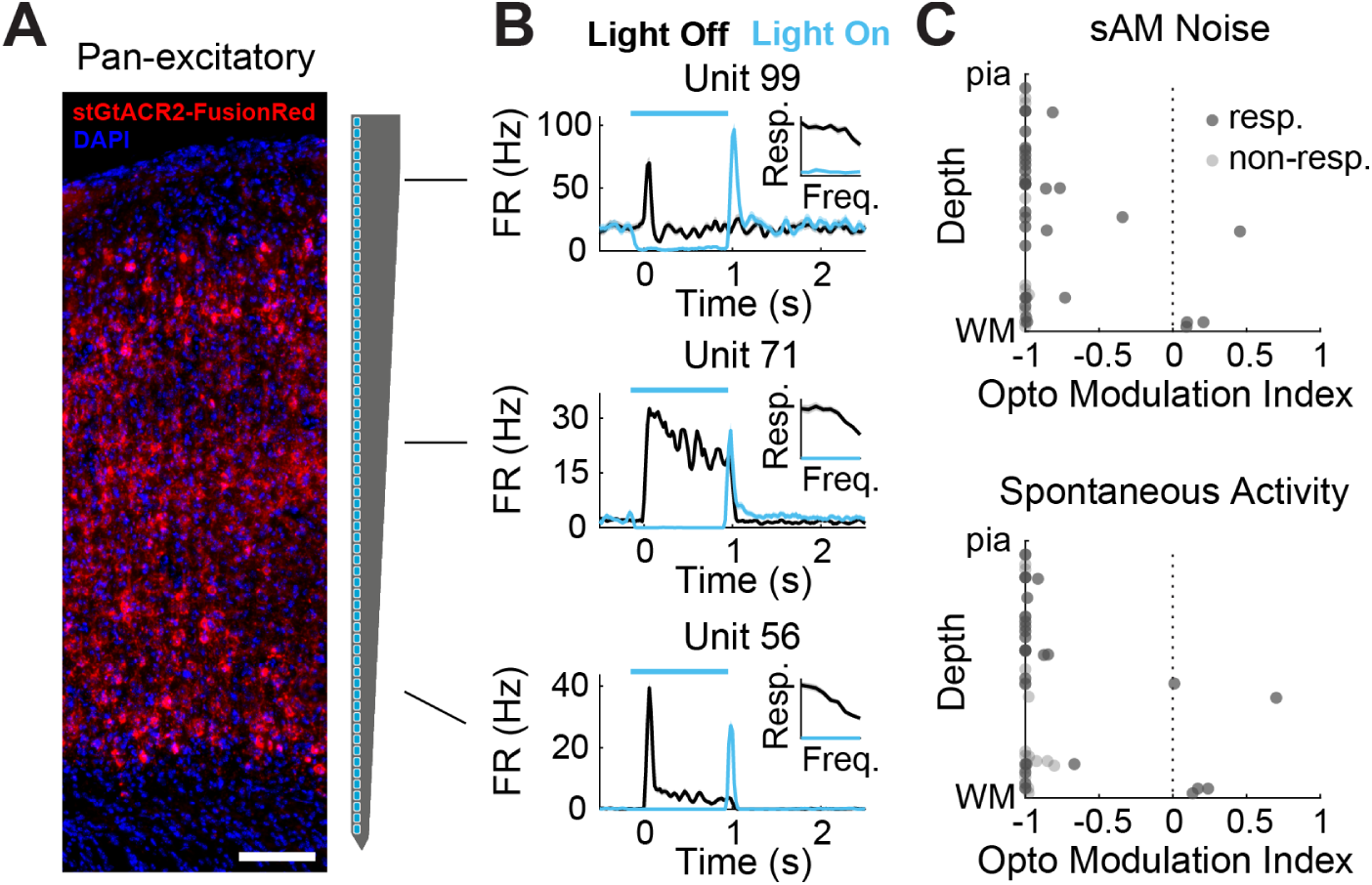
Pan-excitatory optogenetic inhibition of ACtx. **A**: Excitatory neurons in all layers of ACtx of wild-type mice express stGtACR2-FusionRed under the CAMKII promoter (left). Schematic of the 64-channel linear probe used for extracellular recordings (Cambridge NeuroTech, right). **B**: Peristimulus time histograms of exemplar single-unit responses to sAM noise during light off (black) and light on (blue) trials. Insets: Mean responses to each sAM rate. **C**: Optogenetic modulation index as a function of normalized cortical depth for both stimulus-evoked activity (top) and spontaneous activity (bottom) in both sound-responsive (dark gray) and non-responsive (light gray) units.

Optogenetic inhibition of the ACtx significantly impaired the performance of expert mice (N=7), reducing their accuracy from 82.3±0.9% to 59.5±3.5% (**Figure 3A**, paired t-test, *p* = 4.92×10^−4^). To control for the impact of light, a separate cohort of mice (N=4), implanted with optic fibers but lacking a viral injection, were also tested. This group did not show a significant difference in accuracy between light-off and light-on trials (paired t-test: *p* = 0.611), indicating that the observed decrease in performance is specifically attributable to optogenetic inhibition of the ACtx. To investigate the effect of ACtx inhibition across different stimuli, we fit psychometric curves [47–49] to the data. These curves characterize the probability of categorizing a sound as “fast” as a function of sAM rate (**Figure 3D-E**, **Supplementary Figure 1A-B**). Optogenetic inhibition of the ACtx led to a shallower psychometric curve with choice probability approaching 50% across all stimuli. This flattening of the psychometric curve also manifested as a significant increase in the overall lapse rate (**Figure 3B**, paired t-test, control: *p* = 0.072, pan-excitatory: *p* = 7.37 × 10^−5^) as well as decreases in both the psychometric slope (**Figure 2C**, paired t-test, control: *p* = 0.295, pan-excitatory: *p* = 8.21 × 10^−4^), and pseudo d’ (**Supplementary Figure 1C**, two-way ANOVA, interaction between sAM rate and light, control: *p* = 0.186, pan-excitatory: *p* = 6.64 × 10^-3^ ), differences that were not present in the control group.

**Figure 3:**
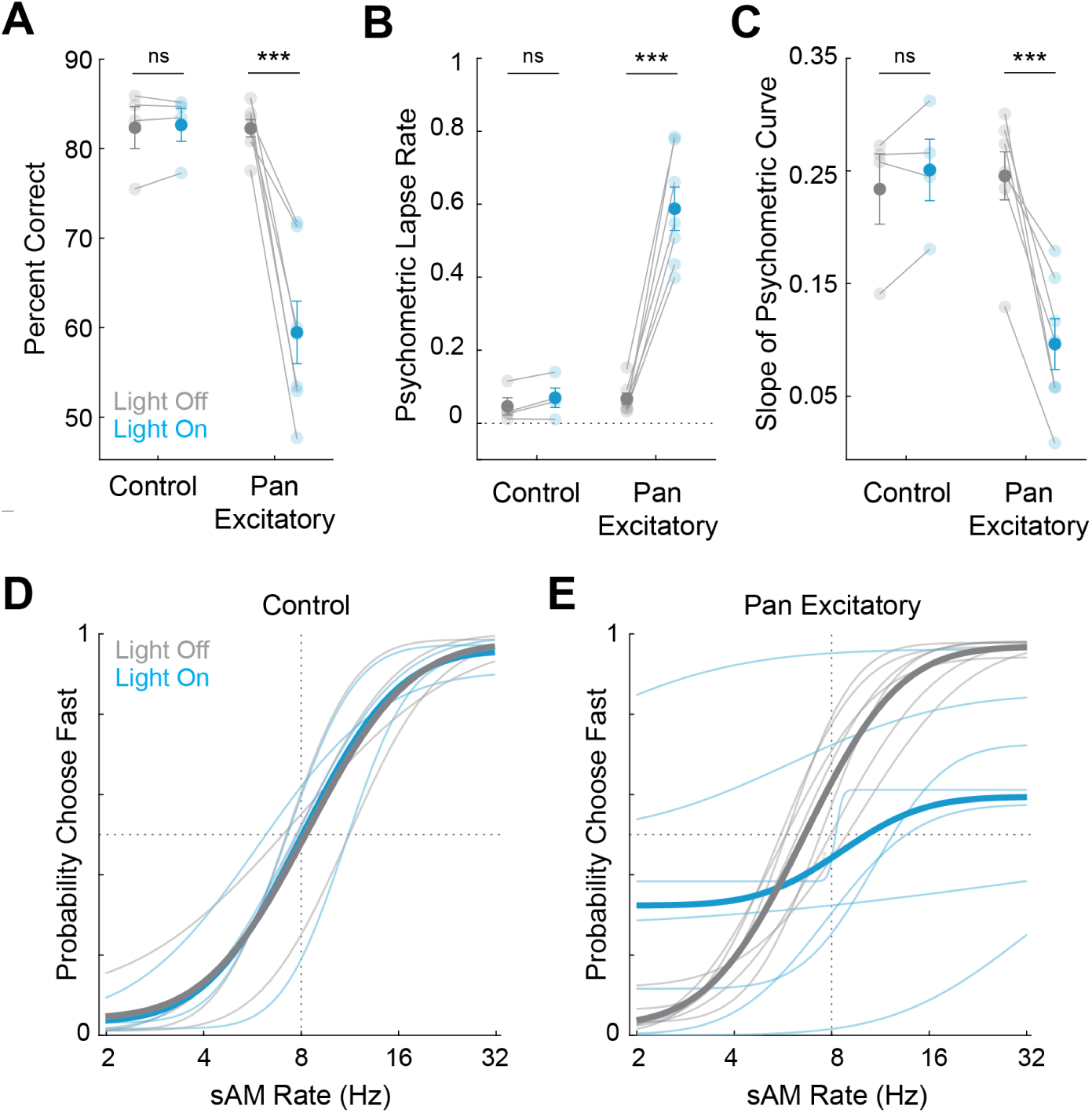
Inhibiting excitatory neurons in ACtx disrupts expression of categorical behavior. **A**: Percent correct for control and pan-excitatory groups, with (blue) and without (grey) light. **B**: Overall lapse rates for control and pan-excitatory groups, with (blue) and without (grey) light. **C**: Psychometric slope for control and pan-excitatory groups, with and without light. **D**: Psychometric curves from control animals with (blue) and without (grey) light. Lighter shades represent individual animals, and bold represents the mean. **E**: Psychometric curves with pan-excitatory suppression (same color conventions as **D**. Asterisks denote statistically significant differences determined using paired t-tests.

Although the average psychometric curve across the population flattened towards chance levels, the psychometric curves exhibited variability across different mice and different sessions. We frequently observed that inhibition drove mice to categorize all sounds uniformly as either “slow” or “fast”, without a consistent bias towards a specific spout or sound association (**Figure 4A**).

**Figure 4:**
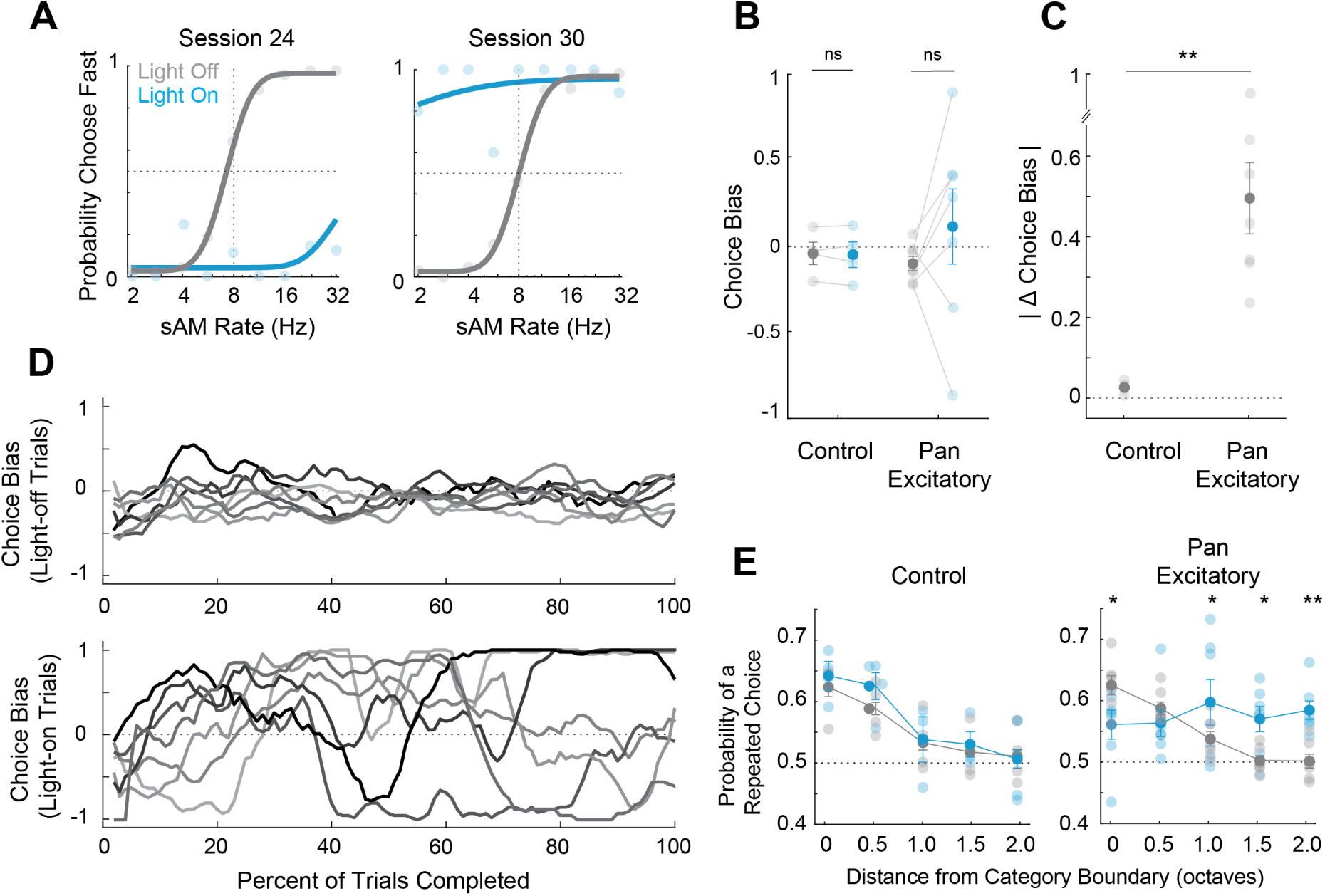
ACtx inhibition leads to a fluctuating choice bias. **A**: Exemplar psychometric curves from two sessions from the same mouse showing a switch in bias from “slow” to “fast”. **B**: Difference in bias with (blue) and without (grey) light. **C**: Difference in absolute bias between light-on and light-off conditions. **D**: Choice bias (computed over a 200 trial sliding window) expressed as a function of learning for pan-excitatory animals in light-off (top) and light-on (bottom) trials. **E**: Proportion of repeated choices as a function of distance from category boundary (octaves). Asterisks denote statistically significant differences determined using an unpaired t-test (C), and a two-way ANOVA (E).

To further investigate this phenomenon, we utilized an asymmetry index to quantify directional choice bias by computing the difference between left and right choices, normalized by the sum of all choices. This measure yields a value whose distance from 0 indicates either left-preferring (*>*0) or right-preferring (*<*0). Considering that half of all stimuli were associated with each spout, we expected expert mice to exhibit a bias close to 0. This expectation was upheld in control mice and in light-off trials of pan-excitatory mice, where the bias was found to be near 0 (**Figure 4B-C**). However, inhibiting the ACtx led to a significant increase in absolute choice bias in expert mice (**Figure 4C**, unpaired t-test, control vs pan-excitatory: *p* = 0.004), consistent with our observation that individual mice often chose one spout in the absence of ACtx activity. This fluctuating choice bias was also present in error trials alone (**Supplementary Figure 2A-B**, unpaired t-test, control vs pan-excitatory: *p* = 0.006). To further examine the consistency of these biases over sessions, we analyzed the bias as a function of the number of complete trials. This visualization revealed strong choice biases in every mouse that flipped sides across learning (**Figure 4D**).

In visual decision-making, task history has a greater impact on choice when the task is more challenging such that mice are more likely to repeat their previous choice when faced with perceptual uncertainty [50]. We examined this in our task by quantifying the proportion of trials in which mice repeated their previous choice as a function of stimuli distance to the category boundary (the task becomes easier and uncertainty decreases as the distance from the category boundary increases). Consistent with previous findings, we found that light-off trials in both groups exhibited the highest proportion of repeated choices close to the category boundary that decreased with distance (**Figure 4E**, two-way ANOVA, main effect for distance, control: *p* = 2.06 × 10^−6^), pan-excitatory: *p* = 0.008). While there was no effect of light on control animals (interaction between light and distance: *p* = 0.725), pan-excitatory inhibition led to a significant change in the probability of repeated choices as a function of distance (interaction between light and distance: *p* = 1.08 × 10^−4^). Notably, with “hard” sounds, 1 - 2 octaves from the category boundary, inhibition led to significantly more repeated choices compared to light-off controls (Sidak’s multiple comparisons test, 1 octave: *p* = 0.046, 1.5 octaves: *p* = 0.020, 2 octaves: *p* = 0.003), comparable to

the proportion observed at and near the category boundary. We also analyzed the probability of a repeated choice as a function of sAM rate and found that, as expected, the probability increased towards the 8 Hz category boundary (**Supplementary Figure 2C**, two-way ANOVA, main effect for sAM rate, control: *p* = 0.002, pan-excitatory: *p* = 0.024), with no effect of light on control animals, and a flattening across all stimuli in the pan-excitatory animals (interaction between light and distance, control: *p* = 0.904, pan-excitatory: *p* = 0.005). This suggests that mice rely primarily on auditory information when the stimulus is easy, and resort to the use of prior information when the stimulus is ambiguous or ACtx is silenced.

### ACtx involvement remains stable across learning

It is feasible that ACtx involvement in categorical learning could diminish across learning, becoming dispensable and acting as a putative tutor to sub-cortical structures. We therefore analyzed performance as a function of trials for both control (**Figure 5A**) and pan-excitatory (**Figure 5B**) mice to examine the effect of ACtx inhibition over learning. As expected, control performance improved over time with no significant differences between performance on light-on and light-off trials (**Figure 5A**, mixed-effect model, main effect for trials: *p* = 9.32 × 10^−11^, main effect for light: *p* = 0.967). In contrast, with pan-excitatory inhibition, the performance on light-on trials differed significantly from light-off trials (**Figure 5B**, mixed-effect model, main effect for trials, *p* = 3.74 × 10^−9^, main effect for light: p=0.007), remaining relatively stable across learning.

**Figure 5:**
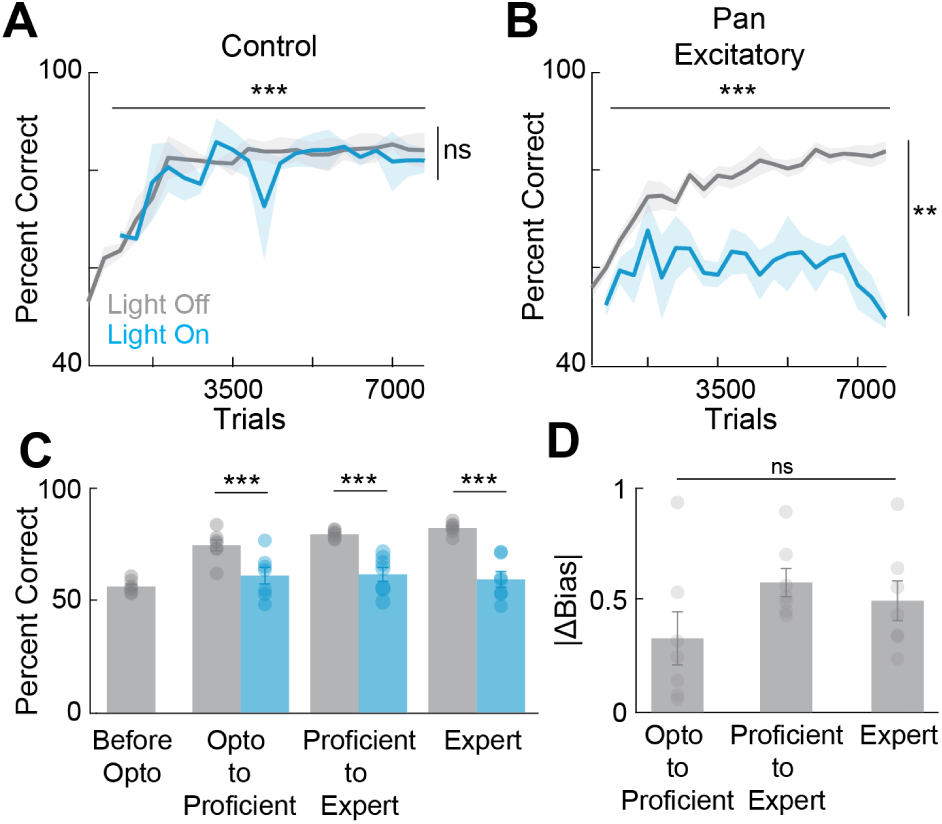
ACtx involvement remains stable across learning. **A**: Percent correct for control mice, with (blue) and without (grey) light, expressed across learning. **B**: Percent correct for pan-excitatory mice, with (blue) and without (grey) light, expressed across learning. **C**: Percent correct for pan-excitatory mice, with (blue) and without (grey) light, computed during four training periods defined by distinct behavioral thresholds. **D**: Change in bias with light for each behavioral training period shown in (C). Asterisks denote statistically significant differences determined using a mixed-effect model (A-B) and a two-way ANOVA (C). Horizontal lines denote main effects across trials, vertical lines denote main effects across light conditions.

To further probe the relationship between the learning and the magnitude of ACtx inhibition, we opted to bin trials across mice based on task performance. This method enabled us to examine the effect of inhibition across mice while controlling for different rates of learning. Trials were grouped into four bins based on distinct performance thresholds: optogenetic introduction (performance *>*60% on easy trials), proficient (performance *>*85% on easy trials) and expert (performance *>*75% on hard trials). This analysis revealed that the contribution of ACtx to sAM categorization remained stable across performance. Early in training, when optogenetic trials were first introduced, ACtx inhibition led to a mean reduction in performance of 13% (**Figure 5C**, two-way ANOVA, Sidak multiple comparisons test, opto to proficient: *p* = 2.26 × 10^−4^). As mice improved their performance on control trials, the effect of ACtx inhibition caused greater reductions in mean performance, from 18% to 23% (**Figure 5C**, two-way ANOVA, Sidak multiple comparisons test, proficient to expert: *p* = 1.53 × 10^−5^, expert: *p* = 1.13 × 10^−6^), commensurate with the increase in performance on light-off trials. Consistent with the influence on performance, we also found that light-induced bias due to inhibition remained stable as a function of learning (**Figure 5D**, one-way ANOVA, main effect for epoch, *p* = 0.08).

### ACtx inhibition effects categorization and not detection

These findings collectively suggest that the ACtx is necessary for the accurate transformation of bottom-up auditory information into perceptual categories. In the absence of ACtx, mice disregard perceptual features when categorizing sounds, instead demonstrating a tendency to lick one spout repetitively. This failure to categorize sounds does not appear to stem from an inability to perceive the sAM rates. Given the unpredictable onset of sound, one would predict that mice would not exhibit reliable licking behavior if they could not perceive the stimulus. However, optogenetic inhibition did not result in a significant change in the rate of no-response trials, during which mice fail to respond within the response window (**Supplementary Figure 3A**, paired t-test, control: *p* = 0.766, pan-excitatory: *p* = 0.293).

Moreover, reaction times for light-on trials in pan-excitatory mice aligned closely with the distribution of light-off trials reaction times across learning (**Figure 6A-B**, two-way anova, main effect for light, opto: *p* = 0.584, proficient: *p* = 0.859, expert: *p* = 0.131). We also observed a notable separation of reaction times to stimuli based on the sAM rates in the later stages of learning, perhaps due to more or less acoustic evidence being necessary to identify “slow” or “fast” rates, respectively (**Figure 6B**, two-way anova, main effect for speed, proficient: *p* = 0.002, expert: *p* = 0.004. The similarity between light-on and light-off trials adds further credence to the notion that mice perceive the differences in sAM rates but fail to accurately categorize them into perceptual groups.

**Figure 6:**
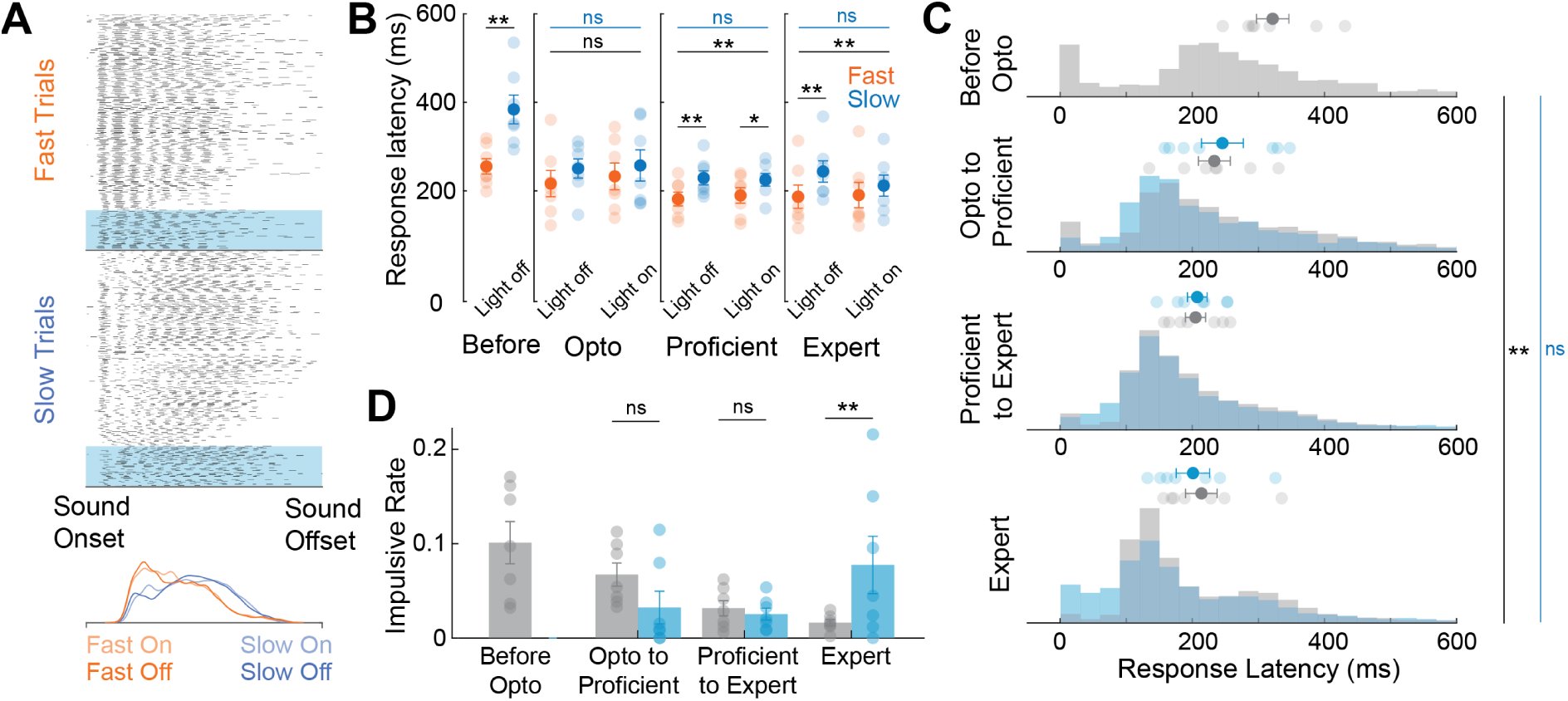
Inhibiting ACtx affects categorization and not detection. **A**: Exemplar lick rasters from one session showing the difference in response latency as a function of sAM rate, with “fast” stimuli (top) associated with shorter latency responses compared to “slow” stimuli (middle) in both light-on and light-off conditions. Probability densities for all groups are shown (bottom). **B**: Average response latencies for “fast” (orange) and “slow” (blue) stimuli, during distinct phases of learning, in both light-off and light-on conditions. **C**: Response latency distributions for light-off (grey) and light-on (blue) trials across learning. **D**: Rate of impulsive licks computed during four distinct training periods. Asterisks denote statistically significant differences determined using a two-way ANOVA (B) and a mixed-effects model (C-D). In B, long horizontal black and blue lines denote main effects for response latency and light, respectively. Short black lines denote multiple comparisons. In C, vertical black and blue lines denote main effects for response latency and light, respectively.

We also examined how the response times evolve as a function of learning by plotting the distribution of response latencies across performance bins for pan-excitatory animals. Response latencies became shorter over the course of training (**Figure 6C**, mixed-effect model, main effect for epoch: *p* = 2.05 × 10^−4^). Inhibition of ACtx led to no significant change in average response latencies (**Figure 6C**, mixed-effect model, main effect for light: *p* = 0.958). However, once mice became experts, the light-on distribution shifted such that more response latencies occurred early after sound onset. Given that the first amplitude modulation of the fastest stimulus occurs 31.25 ms after sound onset, we assumed that any reaction prior to this could not be a result of categorizing the sAM rate. These might instead represent impulsive responses or failure to inhibit anticipatory off-target licking. Early in training (before optogenetic inhibition), impulsive licks were frequent, accounting for ∼10% of all trials (**Figure 6D**). As performance improved, the frequency of impulsive licks decreased in light-off trials (**Figure 6D**, mixed-effect model, main effect for epoch, *p* = 0.005). However, the opposite trend occurred in light-on trials, leading to a significant difference in impulsivity in expert animals (Sidak multiple comparisons test, expert: *p* = 0.006), suggesting that increased impulsivity is related to decreased performance and increased choice bias (**Figure 5C-D**).

### Inhibition of IT and ET neurons impairs the speed and efficacy of categorical learning

The ascending auditory system is critical for the perception and processing of auditory information. Within ACtx, deep-layer pyramidal neurons form major output pathways that propagate information to the rest of the brain. Many of the downstream targets of these projection neurons are essential for decision-making, motor planning, and the execution of auditory-guided behaviors, effectively linking sensation to action [39, 40]. To dissect the specific contributions of ACtx output channels, we extended our pan-excitatory optogenetic manipulations to focus on the three principal classes of excitatory projection neurons. These cell types, intratelencephalic (IT), extratelencephalic (ET), and corticothalamic (CT), represent distinct anatomical and functional streams that transmit cortical output to discrete brain targets. Deep-layer pyramidal neurons exhibit diverse intrinsic and synaptic properties [34], and their axonal projections are organized according to well-conserved subclass-specific rules [32–37]. ITs are found in cortical layers (L) 2/3 - 6, project locally to L2/3, distally to many ipsi- and contralateral cortices as well as the caudal and rostral striatum [51, 52]. They are believed to support intracortical communication and influence motor behavior through corticostriatal pathways [39]. CTs reside in L6a, project locally to L5a, and primarily target the thalamus while collateralizing onto GABAergic cells in the thalamic reticular nucleus (TRN), thereby regulating thalamocortical feedback and sensory gain [53–57]. ETs reside in L5b and target numerous ipsilateral subcortical stations in the midbrain and thalamus, as well as the caudal striatum and amygdala [35, 36]. These corticofugal neurons are increasingly recognized as critical modulators of subcortical plasticity and behavior, particularly during learning [38, 42].

We selectively expressed stGtACR2 in populations of either intratelencephalic (IT), extratelencephalic (ET), or corticothalamic (CT) neurons using a combination of cell-type specific transgenic mice and intersectional viral techniques (**Figure 7A**). To target IT neurons, we used the Tlx3-Cre transgenic mouse line, which provides robust and selective access to L5 IT neurons in the neocortex [58]. For CT neurons, we employed the Ntsr1-Cre line, which reliably labels CT neurons; notably, 97% of Ntsr1-positive neurons in the ACtx are CT, and 90% of all CT neurons are Ntsr1-positive [53]). As transgenic mouse line provides adequate access to ACtx ET neurons, we utilized an intersectional viral strategy that leverages the characteristic projection pattern of ET neurons, their predominant axonal targeting of the inferior colliculus [35–37].

**Figure 7:**
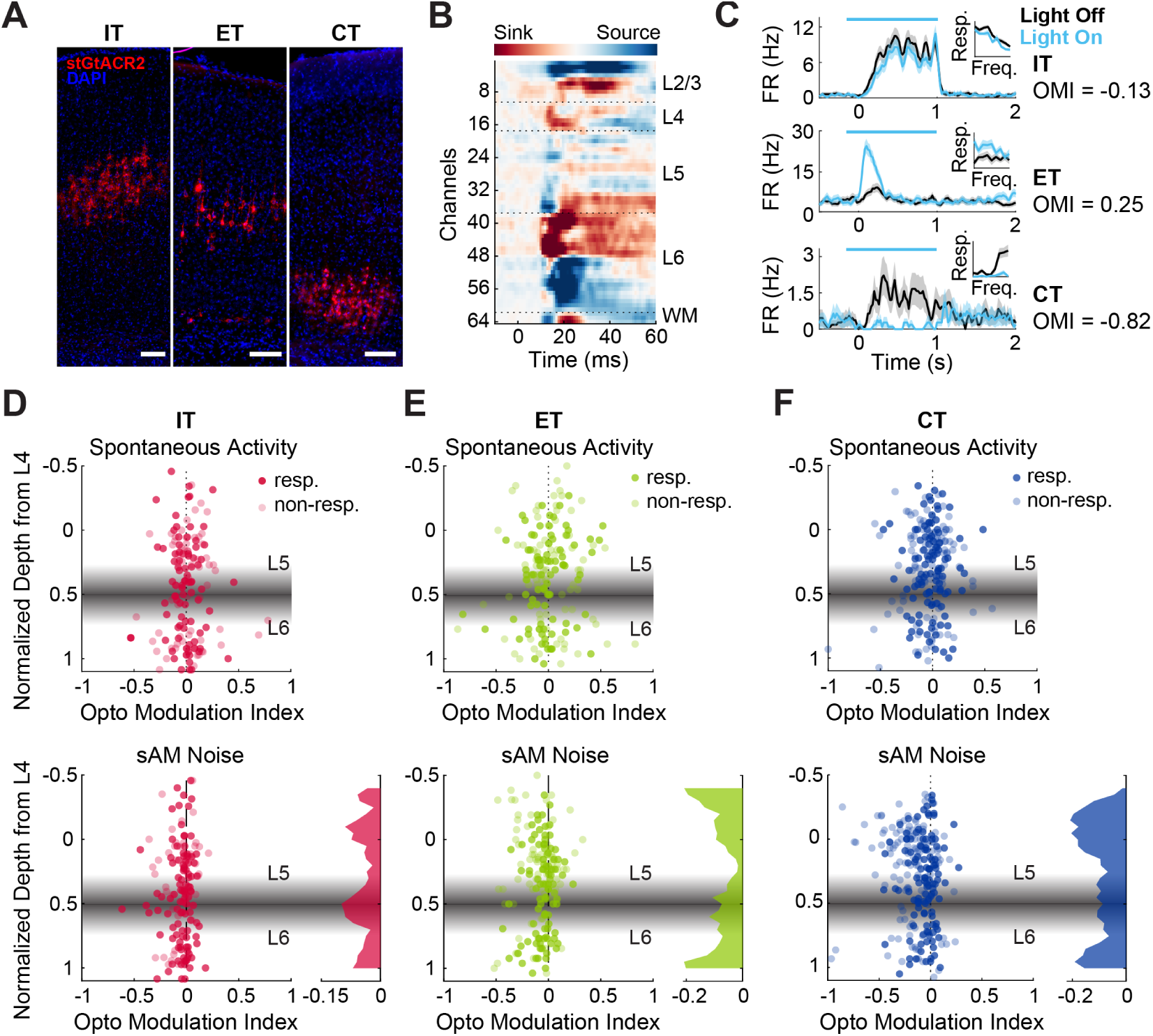
Optogenetic inhibition of distinct ACtx excitatory cell-types. **A**: Cell-type specific expression of stGtACR2-FusionRed in IT (left), ET (middle), and CT (right) neurons in representative ACtx coronal sections. All scale bars 100 *µ*m..**B**: Current source density (CSD) analysis in an example recording delineates laminar boundaries. **C**: Peristimulus time histograms of example single-unit responses to sAM noise during light off (black) and light on (blue) trials. Horizontal blue bars denote the light stimulation epoch. Insets: Mean responses to each sAM frequency. **D**: OMI for all units as a function of normalized cortical depth during suppression of IT neurons spontaneous activity (top) and sound-evoked activity (bottom), in both sound-responsive (dark pink circles) and non-responsive (light pink circles) units. Shaded areas indicate the approximate depth of L5/L6 boundaries. **E**: Same as in (D) for suppression of ET neurons, sound-responsive (dark green circles), non-responsive (light green circles). **F**: Same as in (D) for suppression of CT neurons, sound-responsive (dark blue circles), non-responsive (light blue circles).

We first sought to verify the efficacy of our cell-type-specific optogenetic silencing approach through high-density extraceulluar recordings (IT: N=1, ET: N=1, CT: N=2). Using current source density (CSD) analysis of laminar response profiles to auditory stimuli, we identified the L4 boundary and assigned each recorded neuron to a putative cortical layer (**Figure 7B**). Cell-type-specific activation of stGtACR2 during sound presentation produced heterogeneous effects on cortical activity (**Figure 7C**), consistent with prior findings that selective perturbation of defined excitatory populations can modulate not only local but also translaminar firing and sensory tuning throughout the cortical column [53, 54, 59]. Although the predominant effect was suppression of firing rate (**Figure 7C**, top and bottom), we observed a continuum of modulation across layers and cell-types, including cases of paradoxical enhancement (**Figure 7C**, middle).

To quantify these effects, we computed an optogenetic modulation index (OMI), which compares firing rates during LED-on versus LED-off trials (Figures 7D–F). Silencing L5 IT neurons led to *>*20% suppression in 38 out of 152 recorded units (25%), and *>*20% enhancement in 6 units (4%). A comparable pattern was observed for ET neurons, with 54 of 168 units (32%) showing *>*20% suppression and 10 units (6%) showing *>*20% enhancement. Notably, L6 CT neuron silencing yielded a greater proportion of modulated units, with 117 of 240 units (49%) showing *>*20% suppression and 10 units (11%) showing *>*20% enhancement. This elevated sensitivity to L6 CT silencing is consistent with the known role of L6 CT neurons in dynamically regulating gain and cortical excitability via corticothalamic and intracortical feedback loops [36, 53–55]. These results support the interpretation that even modest perturbations to individual projection classes can produce measurable changes in network dynamics and auditory encoding across cortical layers.

Separate cohorts of IT, ET, and CT mice then underwent identical behavioral training and optogenetic testing protocols as the previously described pan-excitatory and control cohorts. Notably, while broad suppression of excitatory cortical activity significantly impaired performance in expert mice, cell-type-specific inhibition of IT, ET, or CT neurons did not significantly impair either performance (**Figure 8A-B**, paired t-test, IT: *p* = 0.624, ET: *p* = 0.886, CT: *p* = 0.191), or psychometric parameters (**Supplementary Figure 4A-C**), in expert mice. Despite the lack of a strong effect on task expression in expert animals, we hypothesized that even transient inhibition of specific projection types could interfere with the acquisition of stimulus-response associations and thereby impair learning. To investigate this possibility, we examined learning trajectories by plotting behavioral performance as a function of trial number for each group, for both “easy” and “hard” stimuli (**Figure 8C-D**). This analysis revealed significant heterogeneity in learning rates across cell-types, whereby IT- and ET-inhibited mice exhibited slower improvements in performance relative to controls, whereas CT-inhibited mice learned at rates more closely aligned with the control cohort (mixed-effect model, interaction between trial and cell-type, easy stimuli: *p* = 0.004, hard stimuli: *p* = 0.027).

**Figure 8:**
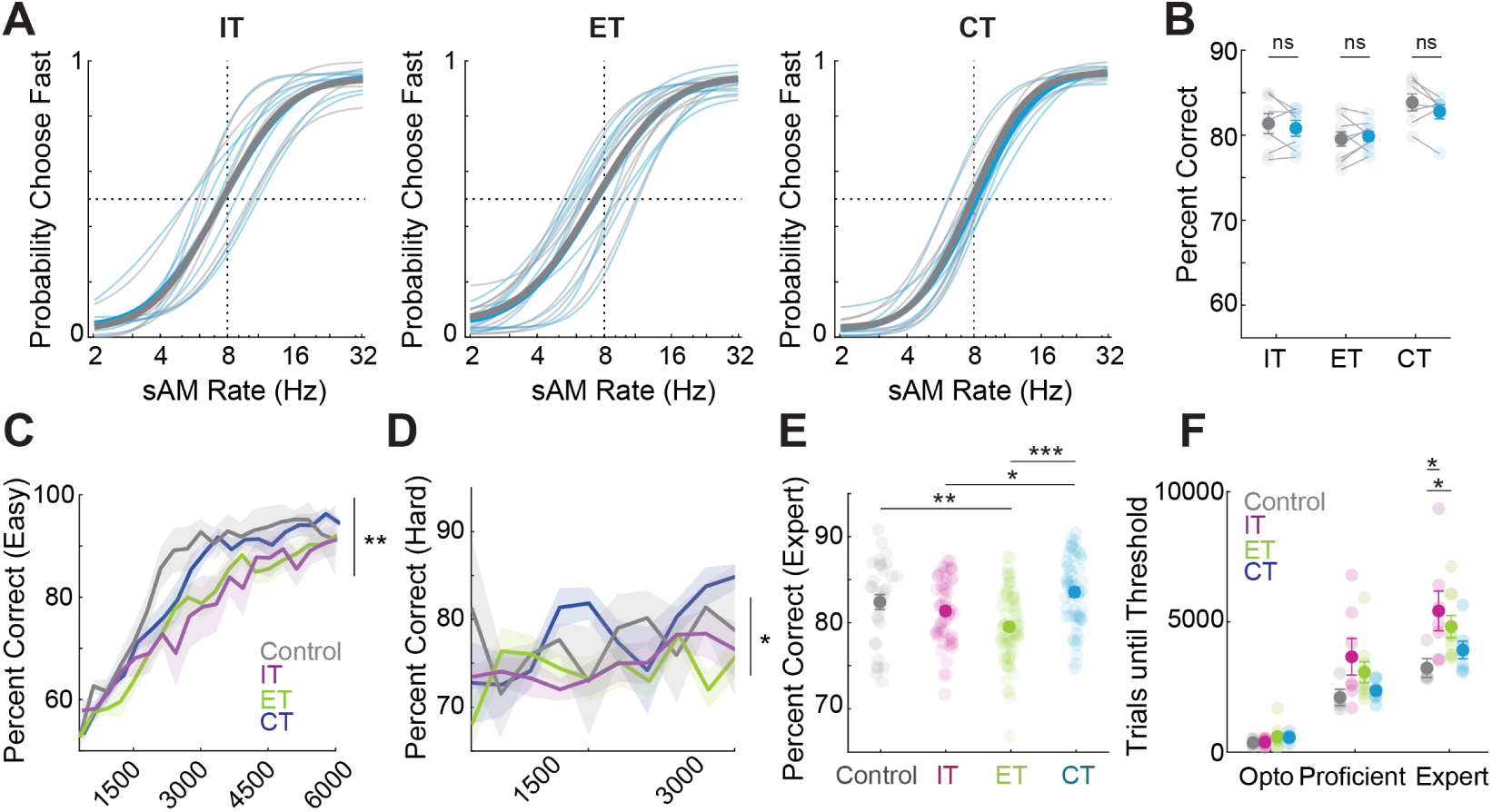
Inhibition of IT and ET neurons impairs speed and efficacy of categorical learning. **A**: Psychometric curves from IT (left), ET (middle), and CT (right) mice for light-off (grey) and light-on (blue) conditions. Lighter shades represent individual animals, and bold represents the mean.**B**: Percent correct for IT, ET, and CT groups, with (blue) and without light (grey). **C**: Percent correct on “easy” trials as a function of trial for control, IT, ET, and CT groups. **D**: Percent correct on “hard” trials as a function of trial for control, IT, ET, and CT groups. **E**: Percent correct for the first eight sessions after reaching expert threshold. **F**: Number of trials for each group to reach optogenetic, proficient, and expert thresholds. Asterisks denote statistically significant differences determined using a mixed-effect model (C-D), one-way ANOVA (E), and a Kruskal-Wallis test (F). Vertical lines denote an interaction between trials and cell-type.

To further quantify learning efficacy and control for variability in within-subject performance across days, we analyzed accuracy across the first eight expert-level training sessions (at which point, performance levels plateau). This revealed a significant reduction in asymptotic performance for ET mice compared to controls and CT-inhibited mice (**Figure 8E**, one-way ANOVA, Tukey’s multiple comparisons test, control versus ET: *p* = 0.007, CT vs ET: *p* = 9.20 × 10^−7^). Additionally, IT-inhibited animals also performed significantly worse than CT-inhibited animals (CT vs IT: *p* = 0.029). These results indicate that perturbation of L5 output pathways, either IT or IT, on as few as 20% of trials, can significantly impede learning. To directly assess learning rate, we calculated the number of trials required to reach our optogenetic, proficient, and expert performance criteria. This analysis revealed that both IT- and ET-inhibited mice required significantly more trials to achieve expert performance relative to controls (**Figure 8F**, Kruskal-Wallis test, *p* = 0.0396, Dunn’s multiple comparisons test, control versus ET: *p* = 0.050, control versus IT: *p* = 0.033).

## Discussion

The role of ACtx in facilitating the acquisition and expression of auditory-guided behaviors is widely reported [17–22, 27]. While these studies have largely focused on whether ACtx is required for the expression of learned behaviors, comparatively little is known about how its causal role may evolve over the course of learning. In particular, the dynamic interplay between cortical plasticity, circuit recruitment, and behavioral strategy adjustment across training remains poorly characterized. In the present study, we address this gap by using a head-fixed auditory choice task paired with transient, trial-specific optogenetic silencing to directly probe how ACtx necessity is modulated as learning progresses. Our results reveal that the contribution of ACtx to categorical behavior is largely static, and remains constant throughout learning. These findings suggest that ACtx plays a dual role: it is essential during initial acquisition, when categorical associations are first being formed, and remains indispensable during later stages, when precise perceptual judgments and optimal strategies are consolidated. ACtx involvement never diminished with training, indicating that even well-learned categorical behaviors continued to rely on ACtx processing. This persistent necessity contrasts with some models of cortical learning that propose a progressive shift from cortical to subcortical control once tasks are automatized. Instead, our findings support a model in which ACtx remains a central hub for transforming acoustic input into categorical decisions even after extensive experience. This view is consistent with recent reports highlighting enduring roles for sensory cortex in expert performance [27, 60].

Transient ACtx silencing had a profound impact on the behavioral strategies employed by mice to perform the categorization task. We observed that optogenetic inhibition introduced a fluctuating choice bias and an increased reliance on prior trial history, suggesting that ACtx is not only critical for processing incoming auditory information but also for integrating sensory evidence to guide adaptive choice behavior. Under conditions of perceptual ambiguity or degraded sensory input, animals often rely on internal priors, such as recent choices or outcomes, to inform current decisions. This is generally considered a hallmark of Bayesian decision strategies [50]. In our task, even high-salience stimuli, which would normally evoke accurate responses, elicited biased, history-dependent behavior during ACtx silencing, implying a generalized failure to extract or interpret sensory evidence. This behavioral pattern mirrors findings from studies implicating posterior parietal cortex in mediating history-dependent decision biases [61]. The fact that ACtx silencing increased history dependence, rather than attenuating it, raises the intriguing possibility that cortical feedback loops between parietal and auditory areas are essential for balancing internal priors against current sensory input.

Analysis of response latencies provides additional insight into the evolving role of ACtx in perceptual decision-making across learning. Late in training, when mice have become proficient at categorizing sounds, optogenetic silencing of ACtx altered the timing of their behavioral responses, increasing impulsive licking. At first glance, these changes might suggest a disruption in the transformation of sensory input into motor output. However, early in learning, when categorization performance is still low, mice nonetheless exhibit robust and timely response latencies, comparable to those in control animals. This implies that basic sensorimotor coupling remains intact even when categorization has not yet developed, and that ACtx silencing does not fundamentally impair motor execution or auditory detection. As learning progresses and animals acquire more refined categorical boundaries, their licking behavior becomes increasingly sensitive to stimulus identity and task demands. We interpret the emergence of faster or more impulsive responses under ACtx inhibition as a sign of impaired stimulus evaluation under conditions of perceptual uncertainty. Without access to cortical processing, mice may default to less informed, and potentially habitual, response strategies in an effort to maintain reward rates. Importantly, the persistence of sound-evoked responses in the absence of ACtx activity, assuming effective and complete silencing, suggests that non-canonical auditory pathways may support rudimentary forms of sensorimotor transformations. One candidate is the thalamo-striatal pathway, which relays auditory information from the thalamus directly to the dorsal striatum, bypassing the cortex entirely [37, 62]. This subcortical circuit has been implicated in driving innate or habitual responses to auditory cues and may provide a residual route for auditory-driven motor actions when cortical processing is compromised.

An important question raised by our findings is whether the observed learning reflects true auditory categorization, or instead simpler forms of behavior such as fine-grained stimulus discrimination or a fixed stimulus–response mapping. Categorization, in its canonical form, requires the generalization of different sensory exemplars into behaviorally meaningful groups based on learned boundaries, rather than simply mapping each stimulus to a unique response. This distinction is critical, as categorization implies the formation of abstract, flexible decision boundaries that can be updated with experience, whereas stimulus-response mapping entails a rigid, itemized association between individual stimuli and motor outputs. Although our current study does not directly manipulate the category boundary, prior work using related auditory paradigms provides compelling evidence that mice and other animals are capable of forming categorical representations of acoustic features. For example, [8] demonstrated that mice trained on a pure tone categorization task could flexibly adapt their behavior when the category boundary was shifted, indicating that animals were not relying on a fixed stimulus-response mapping, but instead had formed a generalized rule or boundary for classifying tones as “low” or “high”. Similarly, work in non-human primates have shown that categorical decisions are supported by neural activity in auditory cortex that reflects category membership rather than physical stimulus features per se [2, 6]. In addition, a key signature of categorical processing is the emergence of sharp decision boundaries, often accompanied by abrupt changes in choice probability around the category boundary, despite smooth physical changes in the stimulus continuum [3]. Our observation that mice exhibit steep psychometric slopes around the category boundary supports the notion that animals are indeed engaging in categorical judgments. While our experiments do not include an explicit category boundary reversal or remapping, our findings are consistent with a growing literature showing that rodents are capable of abstract category learning in the auditory domain. Future experiments involving dynamic boundary shifts or generalization to novel stimuli will be essential to further confirm and dissect the categorical nature of the representations formed during this task.

Although recent advances in genetic and circuit-dissection techniques have begun to reveal the distinct functional roles of distinct excitatory cell-types [35, 38–45], the necessity of these cell-types for behavior, particularly in the context of acquisition versus expression, remains poorly understood. In our study, transient silencing of IT and ET neurons during expert behavior produced subtle but consistent reductions in task expression, while CT neuron silencing had minimal impact. These results suggest that the behavioral contribution of specific projection classes may be graded or context-dependent, rather than absolutely required for task execution under all conditions. When we examined the role of these projection classes in learning, rather than performance alone, marked differences emerged. Mice with targeted inhibition of IT or ET neurons exhibited significantly slower learning trajectories compared to controls, requiring more trials to reach expert performance criteria. In contrast, CT-inhibited mice learned at rates nearly indistinguishable from the control group. These findings indicate that IT and ET neurons are more consequential for the acquisition of sound categories. Our results are consistent with a growing literature implicating ET neurons in driving experience-dependent plasticity and facilitating learning across sensory systems. In the auditory domain, ET neurons have been shown to shape stimulus representations and mediate top-down plasticity during perceptual training [35, 38, 63]. Similar roles for ET neurons have been identified in visual and motor cortices, where they control learning rate and the expression of adaptive cortical dynamics [42, 64]. The present findings suggest that even when a cell type is not strictly necessary for behavioral expression, its transient perturbation during learning can impose lasting constraints on circuit plasticity. An additional interpretation is that our categorization task may not fully engage the descending pathways. Corticofugal projections are thought to be most essential under conditions of perceptual ambiguity, sensory-motor conflict, or when flexible strategy deployment is required [65]. Thus, more challenging tasks or environments that demand behavioral flexibility may more strongly reveal the necessity of deep-layer projection neurons for both learning and decision-making.

Several important methodological considerations must be acknowledged when interpreting the cell-type-specific effects observed in this study. First, silencing was transient, limited to brief windows within trials, and was applied on only 20% of trials. This sparse perturbation paradigm was intentionally designed to minimize long-term compensatory effects and to probe cortical contributions during learning without inducing global circuit dysfunction. However, such an approach may underestimate the full behavioral relevance of each projection class. Second, mirroring our own electrophysiological findings, recent studies have highlighted that silencing individual cell-types (IT, ET, and CT) can profoundly disrupt activity throughout the entire cortical column [53, 59], affecting both ascending and descending signaling streams. These effects underscore the challenge of distinguishing cell-type specificity from population-scale recruitment. Third, while our findings suggest functional specialization across excitatory cell classes, it remains possible that the observed differences are better explained by the number of neurons perturbed rather than their projection identity. This interpretation is supported by our pan-excitatory silencing experiments, in which widespread cortical inhibition had a robust and immediate effect on behavior, likely due to the large number of neurons suppressed across all layers and cell types. Based on single-cell transcriptomic and anatomical reconstructions, ET neurons represent roughly 18% of all excitatory neurons in ACtx, with IT and CT neurons comprising approximately 65% and 17%, respectively [34, 37, 66, 67]. Within L5, IT neurons outnumber ET neurons by roughly 3:1. Yet, our intersectional viral strategy, which restricts opsin expression to retrogradely labeled ET neurons projecting to the midbrain, likely transduced fewer absolute neurons than either the IT or CT (where transgenic mice were used). Thus, the magnitude of the behavioral learning deficit observed with ET silencing cannot be explained by cell number alone, and instead supports the idea that ET neurons may exert a disproportionately large influence on plasticity and behavioral acquisition. Altogether, these methodological factors highlight the need for caution in over-interpreting pathway specificity based on relative behavioral impairments. It remains plausible that all three projection classes contribute meaningfully to behavior, and that the severity of disruption depends not only on cell identity but also on circuit position, output targets, and recruitment dynamics. Future work employing graded silencing approaches, cell-type-specific calcium imaging, and quantitative modeling of circuit output may help disentangle these factors and more precisely determine the relationship between projection class, network dynamics, and behavior.

Ultimately, everything is connected. Cortical contributions to categorization are likely shaped not only by the intrinsic properties of local excitatory and inhibitory circuits, but also by top-down feedback from higher-order regions such as the frontal and parietal cortices, which are known to modulate sensory processing based on task demands, attentional state, and prior expectations. In addition, local inhibitory microcircuits play critical roles in shaping receptive field properties and response gain, thereby influencing how sensory evidence is encoded and read out for decision-making. These processes are further modulated by neuromodulatory systems, including cholinergic, noradrenergic, and dopaminergic inputs, which regulate cortical plasticity, arousal, and behavioral flexibility. Understanding how these diverse influences converge on ACtx to support categorical learning remains a major challenge and provides many distinct avenues for future work.

## Materials and Methods

### Mice

All procedures were approved by the University of Pittsburgh Animal Care and Use Committee and followed the guidelines established by the National Institute of Health for the care and use of laboratory animals. This study is based on data from 40 mice (aged 6-8 weeks, both male and female). All mice were maintained under a 12 hr/12 hr periodic light cycle. 23 C57/Bl6 mice (#000664, Jackson Labs) were used for control groups, as well as viral access to excitatory cells and ET-type cells, 8 Tlx3-Cre mice (B6.FVB(Cg)-Tg(Tlx3-Cre)PL56Gsat/Mmucd, MMRRC) were used to provide genetic access to IT-type cells, and 8 Ntsr1-Cre mice (B6.FVB(Cg)-Tg(Ntsr1-Cre)- GN220Gsat/Mmucd, MMRRC) were used to provide genetic access to CT-type cells.

### Surgical Procedures

#### Virus-mediated gene delivery

Mice were anesthetized using isoflurane (4%) and placed in a stereotaxic plane (Kopf model 1900). A surgical plane of anesthesia was maintained throughout the procedure using continuous infusion of isoflurane (1-2%) in oxygen. Mice lay atop a homeothermic blanket system (Fine Science Tools) that maintained core body temperature at approximately 36.5^◦^C. The surgical area was shaved and cleaned with iodine and ethanol before being numbed with lidocaine (5 mg/ml). For viral delivery to both ACtx, incisions were made to the left and ride side of the scalp to expose the skull around the caudal end of the temporal ridge. The temporalis muscle was then retracted and two burr holes on either hemisphere (approximately 0.3 mm each) were made along each temporal ridge, spanning a region of 1.5-2.5 mm rostral to the lambdoid suture. A motorized stereotaxic injection system (Nanoject III, Drummond Scientific) was used to inject 300 nl of either a non-specific GtACR2 (pAAV1-CamKIIa-stGtACR2-FusionRed, Addgene #105669, titer: 7 × 10^12^ gc/ml), or a Cre-dependent GtACR2 (pAAV1-hSyn1-SIO-stGtACR2-FusionRed, Addgene #105677, titer: 7 × 10 gc/ml) into each burr hole approximately 0.5 mm below the pial surface with an injection rate of 10- 20 nl/min. These parameters yielded a relatively consistent rostro-caudal viral spread across mice of ∼ 2 mm (**Supplementary Figure 5**). Following the injections, the surgical areas were sutured shut, antibiotic ointment was applied to the wound margin, and an analgesic was administered (Carprofen, 5 mg/kg). Mice had ad libitum access to a carprofen-laced medigel (Clear H^2^O) for three days post surgery.

#### Implantation of headplate and optic fibers

Mice were brought to a surgical plane of anesthesia, using the same protocol for anesthesia and body temperature described above. The dorsal surface of the skull was exposed and the periosteum was removed. The skull was first prepared with 70% ethanol and etchant (C&B Metabond) before attaching a custom titanium head plate (eMachineShop) to the skull overlying bregma with opaque dental cement (C&B Metabond). A small craniotomy (0.5 mm x 0.5 mm, medal-lateral x rostral-caudal) was made bilaterally and a ferrule stub (Thorlabs) was implanted over each ACtx surface. Once dry, the cement surrounding the fiber implant was pained black with nail polish to prevent light from escaping. In a subset of mice, a clear-skull preparation was used in lieu of fiber implantation. Here, a thin layer of cyanoacrolite glue was applied to the dorsal surface of the skull, and polished to render the skull optically transparent. Light delivery method did not impact our behavioral results, so all mice were included in our analysis.

### Acoustic and optogenetic stimulation

#### Acoustic stimulation

Stimuli were generated with a 24-bit digital-to-analog converted (National Instruments model PXI- 4461) using custom scripts programmed in MATLAB (Mathworks) and LabVIEW (National Instruments). Stimuli were presented via a free-field speaker (PUI Audio) facing the left (contralateral) ear. Stimuli were calibrated using a wide-band ultrasonic acoustic sensor (SPM0204UD5, Knowles Acoustics).

#### Light delivery

Blue light (488 nm) was generated by either an LED (470 nm, M470F3, Thorlabs) or a diode laser (473 nm, LuxX, Omicron) and delivered bilaterally to the brain via implanted multimode optic fibers coupled to a bifurcating optical patch cable with ceramic mating sleeves. LED/laser power through the optic fiber assembly was calibrated with a photodetector, and set to be ∼2 mW at the fiber tip (Thorlabs).

### Electrophysiology

#### Awake, head-fixed preparation

One to two weeks before acute recording sessions, either a stainless steel ground screw (Grainger) or AgCl wire (World Precision Instruments) was implanted into a small craniotomy on the dorsal surface of the brain and secured with dental cement. Custom titanium headplates were attached as previously described. A small dental cement well was constructed around the right ACtx, leaving the skull intact. Mice recovered for 1-2 days in their home cages with carprofen-laced medigel before acclimatizing to head fixation for 5-7 days. Immediately prior to acute electrophysiological recordings, mice were anesthetized with 2% isofluorane and a small craniotomy (0.5 mm x 0.5 mm, medial-lateral x rostral-caudal) was made over right ACtx using a hand drill with a carbide burr. The well was then filled with cold, sterile saline. Mice recovered from anesthesia while head-fixed in a sound-attenuating booth (Gretch-Ken). At the end of each recording session, the cement chamber was flushed with saline and sealed with a cap of UV-cured cement. Typically, 1-3 recording sessions were performed on each animal over the course of several days. After the final recording, mice were deeply anesthetized with 5% isofluorane and transcardially perfused with cold saline followed by 4% paraformaldehyde (PFA).

#### Data acquisition

At the beginning of each session, a 64-channel, single-shank, silicon probe with 20 µm contact spacing (Cambridge Neurotechnology, ASSY77-H3 or ASSY77-H5) was inserted into the ACtx perpendicular to the pial surface using a micromanipulator (Narishige). Eye ointment was applied over the exposed craniotomy to stabilize the probe and maintain the brain’s moisture. The probe was slowly lowered to a depth of ∼1300 *µ*m, retracted 100-200 *µ*m, then allowed to rest for at least 30 minutes prior to data acquisition. The probe’s reference wire was connected to the implanted screw and kept separate from the ground. Raw signals were amplified, digitized with 16-bit successive approximation, and sampled at 24.4 kHz on a digital headstage (Tucker-Davis Technologies, ZD64 with ZCA-NN64 adapter). Data were acquired, common average referenced, and stored with Synapse (Tucker-Davis Technologies, SI8 & RZ2 BioAmp Processor).

Auditory stimuli were presented pseudorandomly to the left ear of passively listening mice. Sounds lasted for 1 second, followed by a 2 second inter-trial interval. On half of the trials, the brain was illuminated with blue light through a 400 *µ*m fiber (Thorlabs) positioned just above the craniotomy. Light was generated with an LED (as previously described) and calibrated to deliver ∼2 mW at the fiber tip. The light began 100 ms prior to sound onset and lasted for 1.15 s. To assess optogenetic effects on spontaneous activity, trials with no sound were also included. Each stimulus and LED combination was repeated at least 20 times.

#### Current source density analysis

Local field potentials were obtained by low-pass filtering the raw voltage at 300 Hz with a third-order Butterworth filter, downsampling to 1 kHz, and removing significant line noise between 59-61 Hz. Event-related potentials were averaged across trials, and interpolated between channels in 10 *µ*m steps. The CSD was computed as the laplacian approximation of the second spatial derivative, from which the laminar CSD profile could be used to assign a putative layer to each recorded unit.

#### Single-unit analysis

To isolate single-unit activity, the raw data was high-pass filtered (300-10000 Hz) using Kilosort4 [68] with modified spike detection and universal template matching steps to detect only negative-going waveforms. Units were manually inspected with Phy2 (https://github.com/cortex-lab/phy) for waveform quality, refractory period violations, and over-clustering. The following criteria were used to restrict our analysis to well-isolated units:

- False positive rate *<* 0.1, estimated from refractory period violations in a 1.5 ms window
- Cluster isolation distance [69] *<* 15
- Spontaneous firing ≥ 1 Hz or firing evoked by preferred stimulus ≥ 1 Hz on ≥ 50% of trials

Responses were calculated by aligning spikes to the onset of sound and binning in 20 ms intervals. Trial-aligned responses were baseline-subtracted using the average firing rate from 1.1 - 0.1 s prior to sound onset. Responses were averaged for each stimulus and LED combination to generate peristimulus time histograms (PSTHs). Average PSTHs for light-off and light-on responses were generated by combining all stimulus trials before averaging without baseline subtraction. Optogenetic effects on single unit activity were quantified with an optogenetic modulation index (OMI):

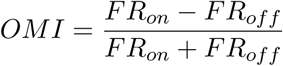

where *FR_on_* and *FR_off_* are the average firing rates during the window from sound onset to LED offset for light on and off trials, respectively. With this index, an OMI of -1 indicates complete suppression. To calculate spontaneous OMI, trials with no sound were analyzed for units with an average spontaneous firing rate ≥ 1 Hz . To evaluate the modulation of sound-evoked responses, units were included with firing rates ≥ 1 Hz for at least one stimulus on ≥ 50% of trials and the OMI was evaluated for each unit’s preferred stimulus. Units were deemed sound-responsive if their PSTH exceeded 3.5 standard deviations above baseline for at least one stimulus condition and responses exceeded this threshold at the same time bin for at least one-third of trials. Since optogenetic modulation was observed in both responsive and non-responsive units, both are included in our analysis. To assess the laminar profile of cell-type specific perturbations, we computed a moving average of OMI values across normalized cortical depth in 0.05 increments with a 0.2 window.

### Mouse Behavior

#### Behavioral training

Approximately a week before training, mice were water restricted to ∼80% of their baseline weight. Mice were first acclimated to handling and head-fixation. After three days of acclimation, mice were introduced to a single spout Pavlovian task in which water reward was dispensed following a 1-second sAM noise stimulus. Once a mouse consistently licked the spout in response to the stimulus, the task was changed to require the mouse to lick in a 1.5 sec response window, 200 ms after sound onset, to trigger the water reward. After the mouse successfully licked for 10 trials in a row in response to sound, a second spout was introduced. Water was dispensed following “fast” sounds (22.8 and 32 Hz) from one spout and “slow” sounds (2 and 2.4 Hz) from the other. To control for potential side biases, the speed/spout association was varied across mice.

Once a mouse was consistently licking both spouts in response to sound, the task was changed to require a lick to the appropriate spout within the response window to trigger the reward. Licks outside of the response window were not counted as a response. Inter-trial intervals were exponentially distributed, and ranged from 4 to 7 s. An incorrect lick triggered a 5 sec timeout before the start of the next trial. Trials with no lick in the response window were counted as “no response” trials with no consequence. Behavioral sessions continued as long as the mouse was engaging in the behavior and sessions were ended when a mouse exhibited persistent “no response” behavior (i.e. *>*10 in a row) suggesting satiation. Once mice had a session with *>*60% correct over 200 trials (light threshold), optogenetic inhibition was randomly introduced on 20% of trials for all following sessions. Once mice reached the threshold of 85% correct across 200 light-off trials (proficient threshold), the remaining sAM noises were introduced (2 - 32 Hz in 0.5 octave steps). A stimulus at the category boundary was included which was not associated with either “fast” or “slow” categories but instead was probabilistically rewarded (50% left/50% right). Mice (N=5) were excluded for failure to appropriately learn the task, defined as *>* 8 sessions to reach the light threshold or *>* 12 sessions to reach the proficient threshold.

#### De-biasing

To prevent the occurrence of spout bias, in which a mouse repeatedly licks one spout regardless of stimulus, we performed a heuristic de-biasing procedure whenever a mouse licked a spout *>*6 times in a row in response to all “easy” stimuli. This de-biasing consisted of repeated presentations of only sounds associated with the non-preferred spout until the mouse correctly responded 3 times in a row. These de-biasing sessions occurred frequently early in learning and tapered off as the animals became proficient at categorization. Optogenetic trials were not included in de-biasing and were not counted towards the criteria for triggering de-biasing. At the beginning of each session, 20-30 trials of “easy” stimuli were presented to ensure unbiased behavior.

#### Behavioral Quantification

Accuracy (percent correct) throughout is quantified as the fraction of correct responses, normalized by the sum of correct and incorrect responses. Response latencies are quantified as the first lick following stimulus onset. Psychometric curves were fit using the following function [47, 48]:

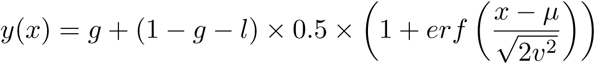

where *y*(*x*) is the probability of a rightward lick and x is the sAM rate and *erf* () is the error function, *g* is the low-frequency lapse rate, *l* is the high-frequency lapse rate (*l*), *µ* is bias (*µ*), and *v* is discrimination sensitivity. Discrimination sensitivity provides the rate of change of performance (psychometric slope). However, estimates of *v* were susceptible to noise due to lapsing, so psychometric slope values were instead computed as the maximum derivative of the fit psychometric function. Bias provides an estimate of the x-axis shift of the psychometric curve, that relates to the current estimate of the category boundary (in our case, 8 Hz), and lapse rates represent “lapses” of performance on easy stimuli, with the overall lapse rate quantified as *g* + *l*. Psychometric error bars represent binomial proportion confidence intervals. A pseudo d’ metric was computed for each stimulus as: *z*(*PC*(*stim_i_*) − *z*(*FAR_right/left_*), where *PC*(*stim_i_*) is the percentage correct for the *i^th^* stimulus, *z* is the inverse of a normal CDF, and *FAR_right_* is the false alarm rate associated with whether the *i^th^*stimulus is on the right of left, *FAR_left_*= *p*(*response_left_*|*stimulus_right_*) and *FAR_right_* = *p*(*response_right_*|*stimulus_left_*).

### Histological Processing and Anatomy

Mice were deeply anesthetized with isoflurane before transcardial perfusion with 4% paraformaldehyde in 0.01 M phosphate buffered saline. Brains were extracted and stored in 4% paraformaldehyde for 12 hr before transferring to cryoprotectant (30% sucrose) until sectioning. Sections (50 *µ*m) were cut using a cryostat (CM1950, Leica), mounted on glass slides, and coverslipped (Vectashield, Vector Laboratories). Fluorescence images were obtained with an epi-fluoresence microscope (Thunder Imager Tissue, Leica) and analyzed using Fiji.

### Statistical Analysis

All statistical analysis was performed with MATLAB (Mathworks) and Prism (GraphPad). Data are reported as mean ± SEM unless otherwise stated. Non-parametric tests were used when data samples did not meet assumptions for parametric statistical tests. In all figures, * (p≤0.05), ** (p≤0.01), and *** (p≤0.001).

## Author Contributions

RFK and RSW conceptualized all experiments. RFK collected and analyzed all data. CNC and MPA conducted all surgical procedures. LIB, JC, RD, HBK, HJM, and JMW assisted with behavioral training. RFK and RSW prepared figures and wrote the manuscript.

## Acknowledgments

We thank current and former members of the Williamson Lab for helpful feedback and discussions and assistance with animal care. Special thanks to Dr. Kameron Clayton for providing comments on an early version of the manuscript. This work was supported by NIH/NIDCD grants R21DC018327 and R01DC020459, a Hearing Health Foundation Emerging Research Grant, and the Klingenstein-Simons Fellowship in Neuroscience to RSW.

---

**Supplementary Figure 1:**
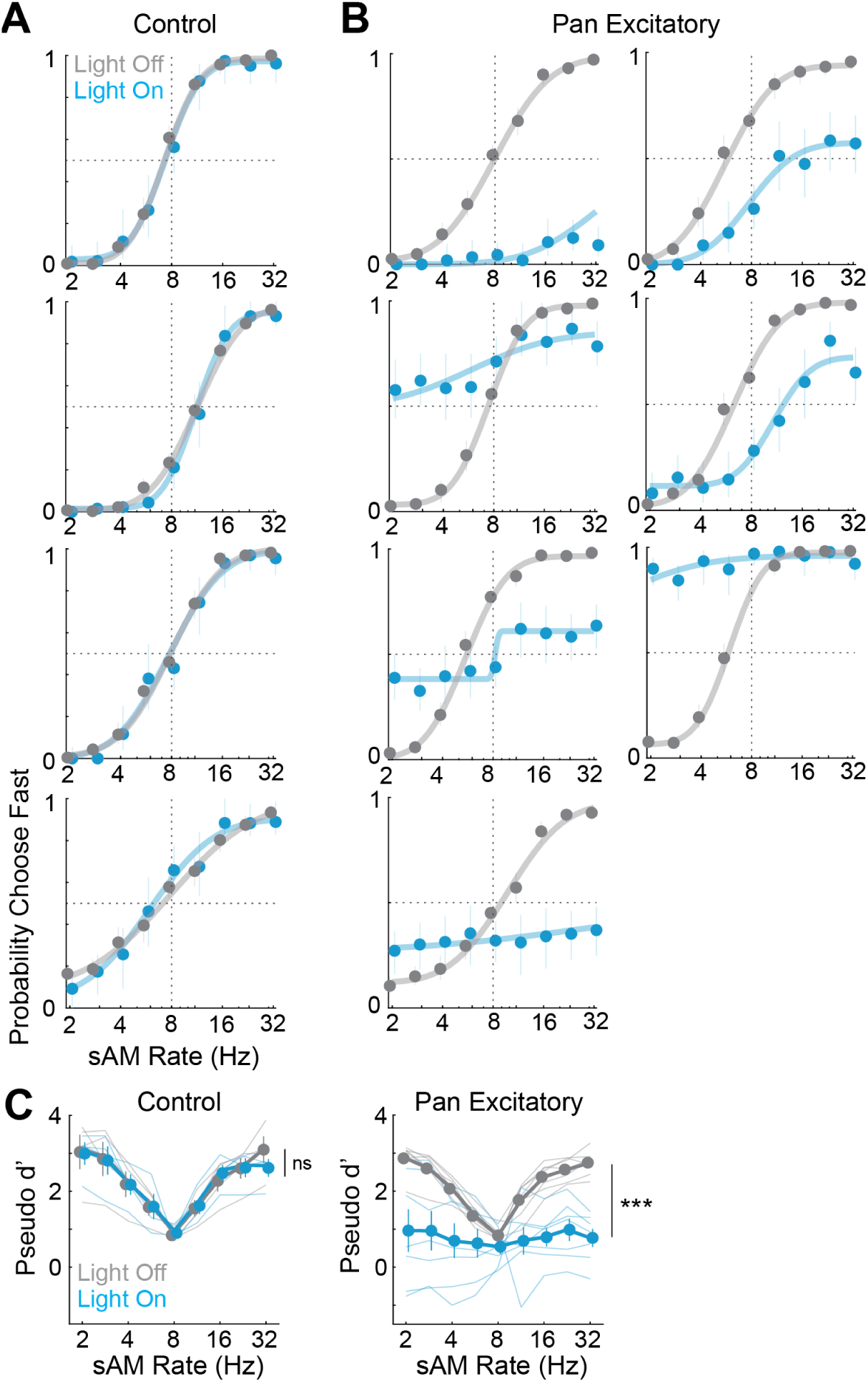
Psychometric curve fits for control and pan-excitatory groups of expert mice. **A**: Psychometric curves from control mice for light-off (grey) and light-on (blue) conditions. Error bars represent binomial proportion confidence intervals. **B**: Psychometric curves from mice with pan-excitatory suppression (same color conventions as **A**. **C**: Pseudo d’ values for all sAM rates, for control (left) and pan-excitatory (right) mice, in light-off (grey) and light-on (blue) conditions. Asterisks denote statistically significant differences determined using a two-way ANOVA. Vertical lines denote a main effect for light.

**Supplementary Figure 2:**
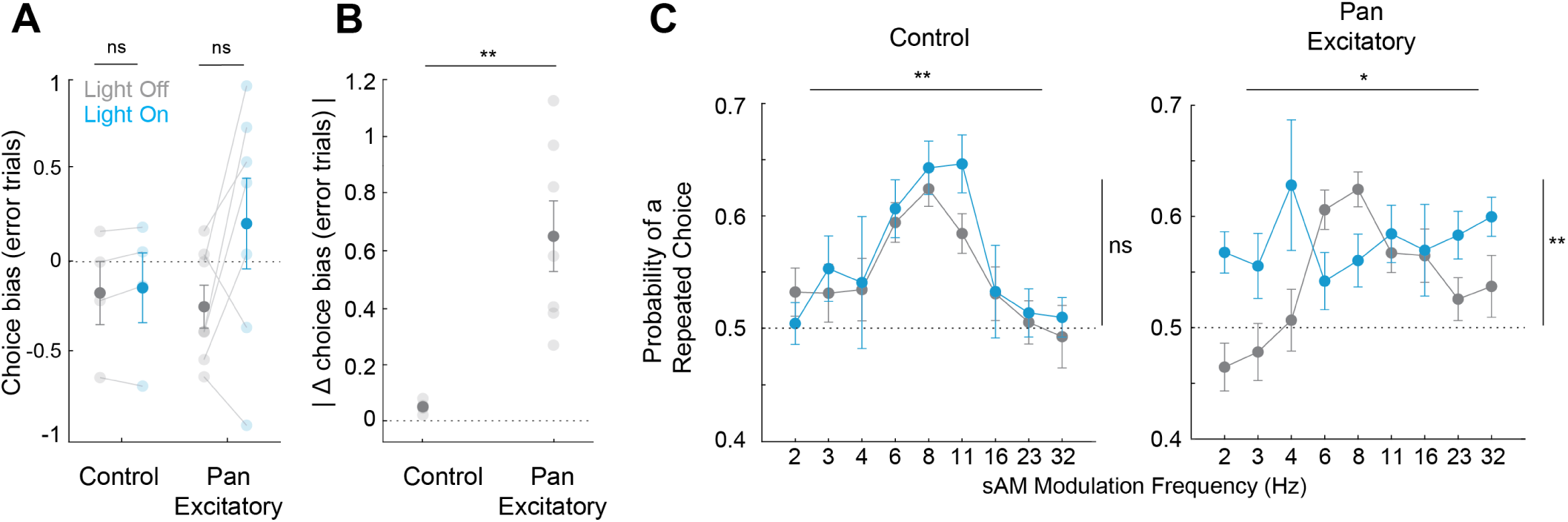
ACtx inhibition leads to a fluctuating choice bias on error trials. **A**: Difference in bias between light-on and light-off conditions in error trials. **B**: Difference in absolute bias between light-on and light-off conditions in error trials. **C**: Proportion of repeated choices as a function of sAM rate (Hz). Asterisks denote statistically significant differences determined using an unpaired t-test (B), and a two-way ANOVA (C). Horizontal lines denote a main effect for sAM rate, and vertical lines denote a main effect for light.

**Supplementary Figure 3:**
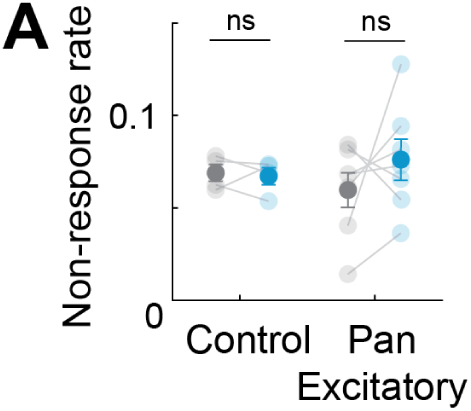
ACtx inhibition does not alter no-response rates. **A**: No-response rate for control and pan-excitatory groups, in light-off (grey) and light-on (blue) conditions.

**Supplementary Figure 4:**
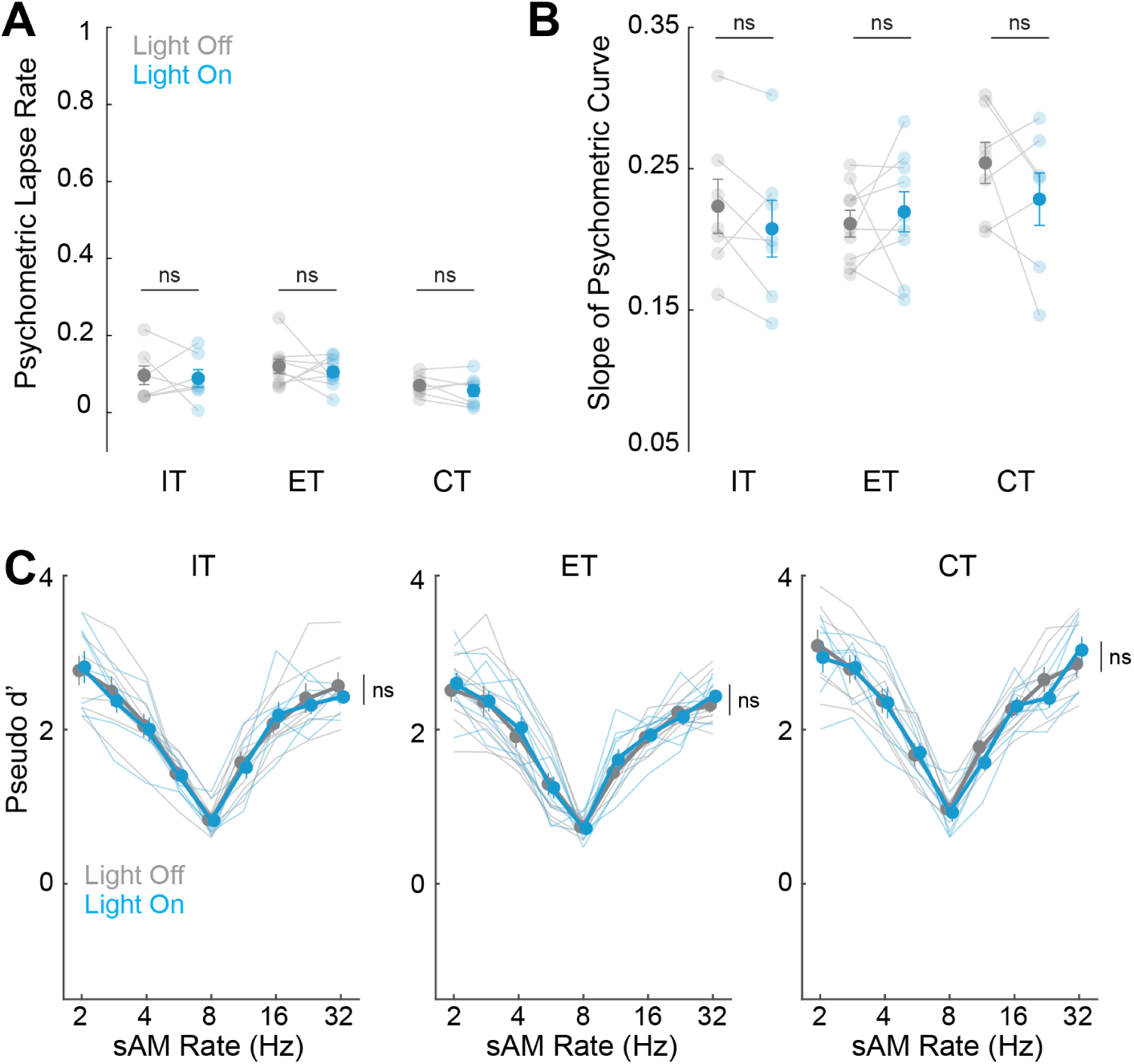
Cell-type specific inhibition does not alter psychometric parameters. **A**: Lapse rate for control and IT/ET/CT groups, with and without light. **B**: Psychometric slope for control and IT/ET/CT groups, with and without light. **C**: Pseudo d’ values for all sAM rates, for IT/ET/CT groups, with and without light.

**Supplementary Figure 5:**
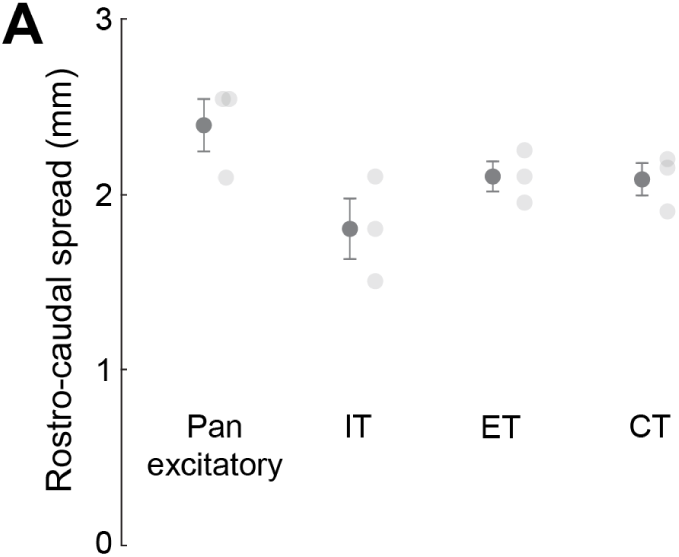
Rostro-caudal spread of stGtACR2. **A**: Rostro-caudal spread of stGtACR2 expression in all groups of mice.

